# Ribovore: ribosomal RNA sequence analysis for GenBank submissions and database curation

**DOI:** 10.1101/2021.02.11.430762

**Authors:** Alejandro A. Schäffer, Richard McVeigh, Barbara Robbertse, Conrad L. Schoch, Anjanette Johnston, Beverly A. Underwood, Ilene Karsch-Mizrachi, Eric P. Nawrocki

**Affiliations:** Cancer Data Science Laboratory, National Cancer Insitute, National Institutes of Health, Bethesda, MD, 20892 USA; National Center for Biotechnology Information, National Library of Medicine, National Institutes of Health, Bethesda, MD, 20894 USA

**Keywords:** ribosomal RNA, annotation, alignment, ncRNA

## Abstract

**Background:** The DNA sequences encoding ribosomal RNA genes (rRNAs) are commonly used as markers to identify species, including in metagenomics samples that may combine many organismal communities. The 16S small subunit ribosomal RNA (SSU rRNA) gene is typically used to identify bacterial and archaeal species. The nuclear 18S SSU rRNA gene, and 28S large subunit (LSU) rRNA gene have been used as DNA barcodes and for phylogenetic studies in different eukaryote taxonomic groups. Because of their popularity, the National Center for Biotechnology Information (NCBI) receives a disproportionate number of rRNA sequence submissions and BLAST queries. These sequences vary in quality, length, origin (nuclear, mitochondria, plastid), and organism source and can represent any region of the ribosomal cistron.

**Results:** To improve the timely verification of quality, origin and loci boundaries, we developed Ribovore, a software package for sequence analysis of rRNA sequences. The `ribotyper` and `ribosensor` programs are used to validate incoming sequences of bacterial and archaeal SSU rRNA. The `ribodbmaker` program is used to create high-quality datasets of rRNAs from different taxonomic groups. Key algorithmic steps include comparing candidate sequences against rRNA sequence profile hidden Markov models (HMMs) and covariance models of rRNA sequence and secondary-structure conservation, as well as other tests. Nine freely available `blastn` rRNA databases created and maintained with Ribovore are used for checking incoming GenBank submissions and used by the `blastn` browser interface at NCBI. Since 2018, Ribovore has been used to analyze more than 50 million prokaryotic SSU rRNA sequences submitted to GenBank, and to select at least 10,435 fungal rRNA RefSeq records from type material of 8,350 taxa.

**Conclusion:** Ribovore combines single-sequence and profile-based methods to improve GenBank processing and analysis of rRNA sequences. It is a standalone, portable, and extensible software package for the alignment, classification and validation of rRNA sequences. Researchers planning on submitting SSU rRNA sequences to GenBank are encouraged to download and use Ribovore to analyze their sequences prior to submission to determine which sequences are likely to be automatically accepted into GenBank.

## Background

In 1977, Carl Woese and George Fox proposed the *Archaebacteria* (later renamed *Archaea*) as a third domain of life distinct from *Bacteria* and *Eukaryota* based on analysis of small subunit ribosomal RNA (SSU rRNA) oligonucleotide fragments from 13 microbes [1]. The use of SSU rRNA to elucidate phylogenetic relationships continued and dramatically expanded in the late 1980s when Norm Pace and colleagues developed a technique to PCR amplify rRNA genes from potentially unculturable microbes in environmental samples by targeting so-called *universal primer sites* [2]. The technique was later refined by Pace and others including Ward, Weller [3] and Giovanonni and colleagues [4]. Environmental studies targeting SSU rRNA as a *phylogenetic marker gene* that seek to characterize the diversity of life in a given environment have remained common ever since, and consequently there are now millions of prokaryotic SSU rRNA sequences in public databases. When rRNA sequences are submitted to public databases, such as GenBank, it is important to do quality control, so that subsequent data analyses are not misled by errors in sequencing and sequence annotation. Because rRNA gene sequences do not code for proteins, but have been studied so extensively, specialized checks for correctness and completeness are feasible and desireable. The focus of this paper is the description of Ribovore, a software package for validating incoming rRNA sequence submissions to GenBank and for curating rRNA sequence collections.

SSU rRNA was initially chosen by Woese and Fox for inferring a universal phylogenetic tree of life because it existed in all cellular life, was large enough to provide enough data (about 1500 nucleotides (nt) in *Bacteria*), and had evolved slowly enough to be comparable across disparate groups [5]. The first environmental surveys targeted SSU rRNA, but studies targeting LSU rRNA, which is roughly twice as long as SSU rRNA, followed soon after [6, 7]. These types of analyses eventually began to target eukaryotes, especially *Fungi* [8].

In eukaryotes, the 5.8S rRNA gene is surrounded by two internal transcribed spacers (ITS1 and ITS2). This region is sometimes collectively referred to as the ITS region and it has been selected as the primary fungal *barcode* since it has the highest probability of successful identification for the broadest range of *Fungi* [9]. However, the LSU rRNA gene [10] is a popular phylogenetic marker in certain fungal groups [9]. In general, the nuclear SSU rRNA has poor species-level resolution in most *Fungi* and other eukaryote taxonomic groups [11, 9], but remains useful at species-level phylogenetic inference in some rapid evolving groups such as the diatoms [12]. Species identification in protists takes a two-step barcoding approach, which use the ~500 bp variable V4 region of the SSU rRNA gene as a variable marker and then use a group-specific barcode for species-level assignments, some of which include the LSU rRNA gene and ITS region [11].

Specialized analysis tools and databases have been developed to help researchers analyze their rRNA sequences. Many of these specialized tools are based on comparing sequences to either profile hidden Markov models (profile HMMs) or covariance models (CMs). CMs are profile stochastic context-free grammars, akin to profile HMMs of sequence conservation [13, 14], with additional complexity to model the conserved secondary structure of an RNA family [15, 16, 17]. Like profile HMMs, CMs are probabilistic models with position-specific scores, determined based on the frequencies of nucleotides at each position of the input *training* alignment used to build the model. Unlike HMMs, CMs also model well-nested secondary structure, provided as a single, fixed consensus secondary structure for each model and annotated in the input training alignment. A CM includes scores for each of the possible 16 (4×4) basepairs for basepaired positions and both paired positions are considered together by scoring algorithms.

The incorporation of secondary structure has been shown to significantly improve remote homology detection of structural RNAs [18], and for SSU rRNA considering structure has been shown to offer a small improvement to alignment accuracy versus profile HMMs [19, 20]. For eukaryotes, where SSU and LSU rRNA sequences are often more divergent at the sequence level than for *Bacteria* and *Archaea*, harnessing structural information during alignment may be more impactful.

The specialized tools for rRNA sequence analysis include: databases, some of which are integrated with software, rRNA prediction software, and multiple alignment software. The integrated and highly curated databases include the ARB workbench software package for rRNA database curation [21], the Comparative RNA Website (CRW) [22], the Ribosomal Database Project (RDP) [23, 24], and the Greengenes [25], and Silva [26] databases. These databases differ in their scope and methodology. CRW contains tens of thousands of sequences and corresponding alignments of SSU, LSU and 5S rRNA from all three domains as well as from organelles, along with secondary structure predictions for selected sequences. Greengenes, which is seemingly no longer maintained as its last update was in 2013, includes SSU rRNA sequences for *Bacteria* and *Archaea*, but not for *Eukarya*, nor does it contain any LSU rRNA sequences. RDP also includes SSU rRNA for *Bacteria* and *Archaea*, as well as fungal LSU rRNA and ITS sequences, but no other LSU sequences. Silva, which split off from the ARB project starting in 2005 [27], includes bacterial, archaeal and eukaryotic (fungal and non-fungal) SSU and LSU rRNA sequences. RDP includes more than 3 million SSU rRNA and 125,000 fungal LSU rRNA sequences as of its latest release (11.5), and Silva includes more than 9 million SSU rRNA and 1 million LSU rRNA sequences (release 138.1).

Available rRNA prediction software packages include RNAmmer [28], rRNAselector [29], and barrnap(https://github.com/tseemann/barrnap) all of which use some version of the profile HMM software HMMER [30] to predict the locations of rRNAs in contigs or whole genomes.

Both RDP and Silva make available multiple alignments of all sequences for each gene and taxonomic domain, and all include several sequence analysis tools for tasks such as classification. The alignment methodology differs: Silva uses SINA, which implements a graph-based alignment algorithm that computes a sequence-only based alignment of an input sequence to one or more similar sequences selected from a fixed reference alignment [31]. RDP uses Infernal [32], which computes alignments using CMs.

Per-domain CMs for SSU and LSU rRNA are freely available in the Rfam database, a collection of more than 3900 RNA families each represented by a consensus secondary structure annotated reference alignment called a *seed* alignment and corresponding CM built from that alignment [33]. Rfam includes five full length SSU and four full length LSU rRNA families and CMs. Although RDP uses CMs for rRNA alignment, the CMs are not from Rfam. Users can download and use Rfam CMs to annotate their own sequences using Infernal, thus offering a distinct strategy from Silva or RDP for rRNA analysis.

The Rfam database includes a model (RF02542) for SSU rRNA from *Microsporidia*, a phylum of particular interest within the kingdom of *Fungi*. More than 30 years ago, Woese and colleagues discovered that *Microsporidia* have a distinctive ribosome that is smaller and more primitive than the ribosomes of most if not all other eukaryotes [34]. Recently, Barandun and colleagues presented the first crystal structure of the ribosome of *Microsporidia*, confirming that both the SSU and LSU rRNA are smaller than in other *Fungi* [35]. Most of the sequence analysis and curation to date in *Microsporidia* has focused on SSU rather than LSU rRNA.

### GenBank processing of rRNA sequences

Data submitted to GenBank are subject to review by NCBI staff to prevent incorrect data from entering NCBI databases. Over the past three decades, personnel called *GenBank indexers* have spent a large proportion of their time validating incoming submissions of thousands to millions of rRNA sequences due to the large number of rRNA sequences generated in phylogenetic and environmental studies. Similarity searches with `blastn` have been used to compare submitted rRNA sequences against databases of trusted, high-quality rRNA sequences for prokaryotic 16S and 23S, eukaryotic 18S and 28S, and chloroplast-specific sequences. However, these databases were not comprehensive before curation with Ribovore began. The blastn query results were a primary source of evidence used to determine if rRNA sequences would be accepted to GenBank or not. Prior to the Ribovore project, suitable blastn databases did not exist for validating submissions of eukaryotic SSU rRNA or LSU rRNA sequences, making checking for those genes especially difficult and time-consuming.

Starting in 2016, a system with predefined criteria for per-sequence blastn results was deployed at NCBI; submissions in which all sequences met those criteria have been automatically accepted into GenBank without any indexer review. Ribovore-based tests began being used in conjunction with or instead of `blastn`-based tests for some submissions in this system in June 2018. Although the engine inside the pre-2018 validation system, BLAST, is freely available and portable, the system as a whole was internal to GenBank and not portable, preventing researchers who wish to submit sequence data to GenBank (henceforth, called “submitters”) from replicating the tests on their local computers.

For rRNA sequences as well as other sequences of high biological interest, GenBank indexers and other NCBI personnel want to carry out two related and recurrent processes: quick identification of which submitted sequences should be accepted into GenBank, and the construction of non-redundant collections of trusted, full length sequences that have no or few errors. The second problem is the motivation behind the entire RefSeq project [36]. Towards addressing the first problem, the development of an alternative sequence validation system for rRNA included four design goals offering potential improvements over the existing system. First, the system should be as deterministic and as reproducible as possible in deciding whether sequences are accepted or not, which we refer to as *passing* (accepted) or *failing* (not accepted), allowing submissions with zero failing sequences to be automatically added to GenBank without the need for any manual GenBank indexer intervention. It is unavoidable that automated decisions about acceptance can change over time because various inputs to the system, such as the NCBI taxonomy tree, change over time. Second, the system should be available as a standalone tool that submitters can run on their sequences prior to submission, saving time for both the GenBank indexers and submitters. Third, the system should be general enough to facilitate extension to additional taxonomic groups and rRNA genes. Fourth, the system should be capable of increasing the stringency of tests for quality and adding tests to avoid redundancy to enable producing collections of high quality non-redundant sequences for other applications, such as serving as blastn databases.

Because none of the existing databases or specialized rRNA tools listed above address all of these design goals, we implemented the freely available and portable Ribovore software package for the analysis of SSU rRNA and LSU rRNA sequences from *Bacteria*, *Archaea*, and *Eukarya* as well as mitochondria from some eukaryotic groups. Ribovore includes several programs designed for related but distinct tasks, each of which has specific rules dictating whether a sequence passes or fails based on deterministic criteria described in detail in the Implementation section and in the Ribovore documentation. `rRNA_sensor` is a simplified, standalone version of the previous `blastn`-based system that is more portable and faster for bacterial and archaeal SSU rRNA than the previous system owing to a smaller blastn target database constructed by removing redundancy from the pre-existing `blastn` database. `ribotyper` is similar to `rRNA_sensor` but compares each input sequence against a library of profile HMMs and/or CMs offering an alternative, and in some cases, more powerful approach than the single sequence-based `blastn` algorithm. Additionally, `ribotyper` can be used to validate the taxonomic domain each sequence belongs to because it compares a set of models from different taxonomic groups against each sequence. To take advantage of both single-sequence and profile-based approaches, and partly to ease the transition from the previous `blastn`-based system towards profile-based analysis, we implemented `ribosensor` that runs both `rRNA_sensor` and `ribotyper` and then combines the results. Up to this point, `rRNA_sensor` and `ribotyper` are deliberately designed to accept both partial and complete sequences of moderate quality or better. To more selectively identify full-length rRNA sequences that extend up to, but not beyond the gene boundaries, we implemented `riboaligner` which runs `ribotyper` as a first pass validation, and then creates multiple alignments and selects sequences that pass based on those alignments. Finally, to make Ribovore capable of generating datasets of trusted sequences from different taxonomic groups for wider use by the community, we developed `ribodbmaker`, which chooses a non-redundant set of high-quality, full-length sequences based on a series of tests. The pipeline of tests includes some specific to rRNA, including analysis by `ribotyper` and `riboaligner`, some more general tests, such as counting ambiguous nucleotides and vector contamination screening, and some tests that require connection to the NCBI taxonomy database to validate the taxonomy assignment of sequences.

## Implementation

Ribovore is written in Perl and available at https://github.com/ncbi/ribovore. The Ribovore installation procedure also installs the program `rRNA_sensor`, which is described here as well. The `rRNA_sensor` program includes a shell script and Perl scripts and is available at https://github.com/aaschaffer/rRNA_sensor. These packages use existing software as listed in Table 1.

**Table 1.**
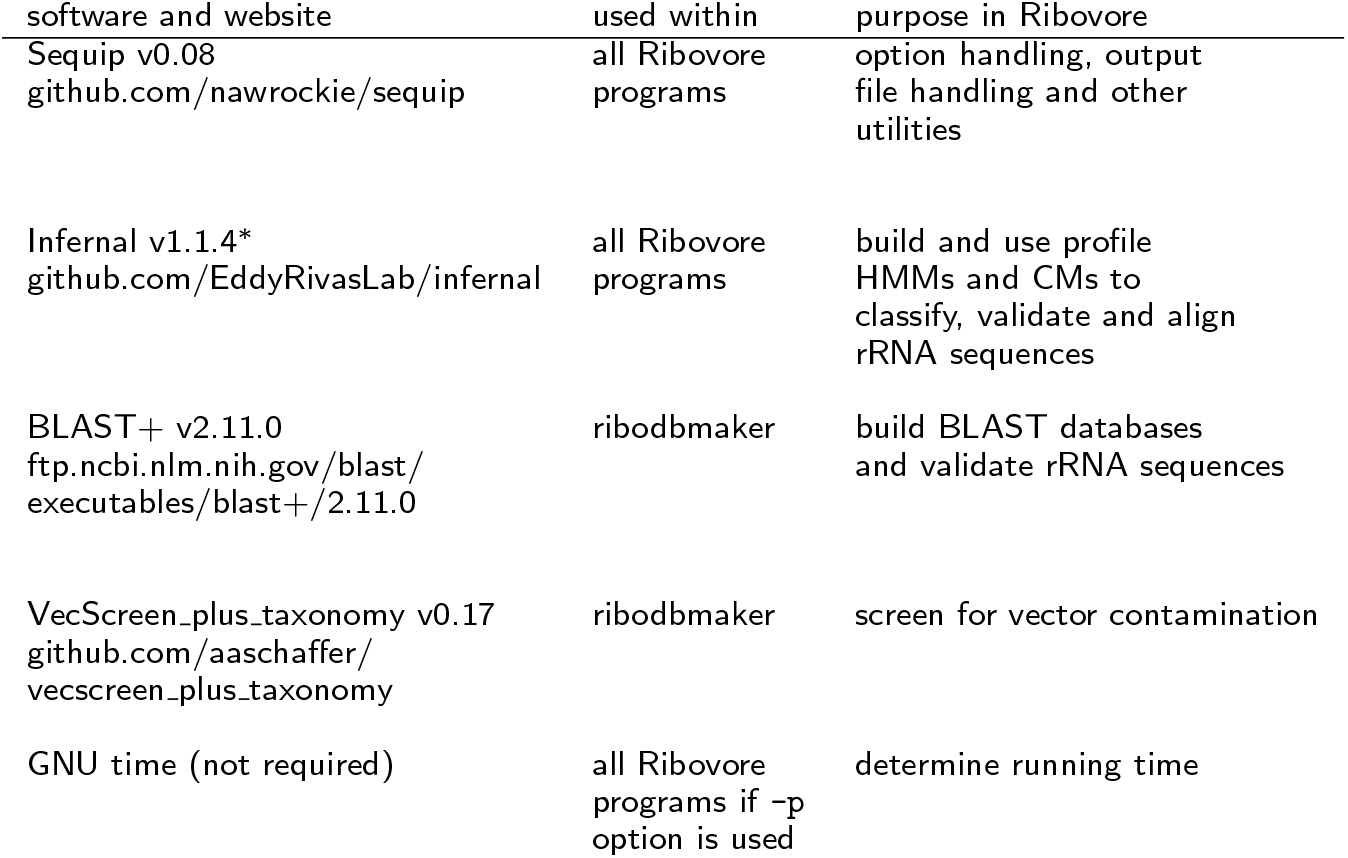
Software packages and libraries used within Ribovore v1.0. *: the esl-cluster executable from Infernal v1.1.2, which is absent in v1.1.4, is also installed and used within Ribovore.

Each of the four Ribovore programs takes as input two command-line arguments: the path to an input sequence file in FASTA format and the name of an output directory to create and store output files in. Command-line options exist to change default parameters and behavior of the programs in various ways. The options as well as example usage can be found as part of the source distribution and on GitHub in the form of markdown files in the Ribovore documentation subdirectory (e.g. https://github.com/ncbi/ribovore/blob/master/documentation/ribotyper.md). Central to each of the scripts is the concept of sequences passing or failing. If a sequence meets specific criteria, many of which are changeable with command-line options, then it will pass and otherwise it will fail, as discussed more below. An overview of the four Ribovore programs and `rRNA_sensor` is shown in Figure 1.

**Figure 1.**
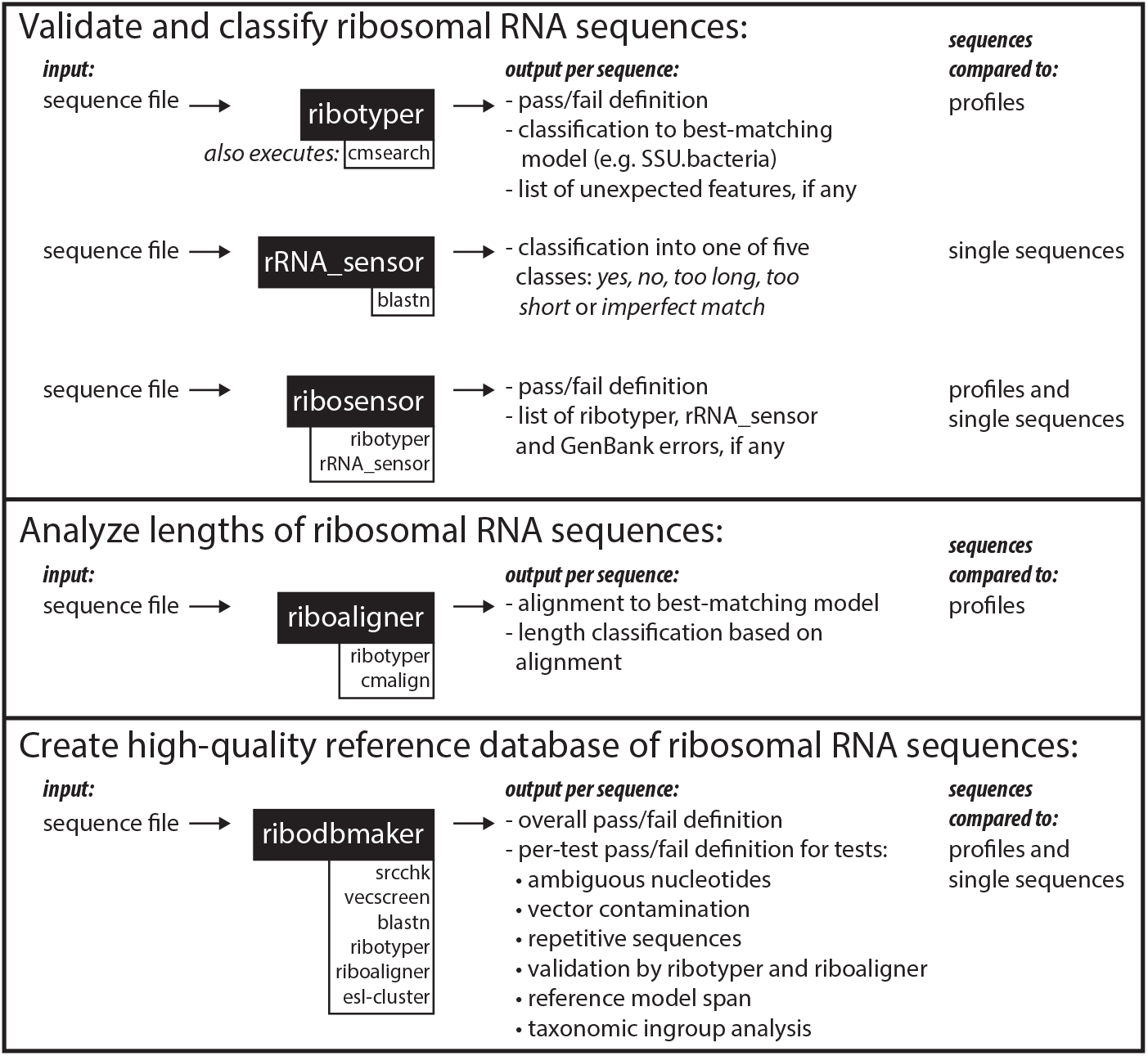
Schematic summarizing the use cases for the four Ribovore programs and rRNA_sensor. Programs listed in white boxes underneath the black boxes are important external programs executed from within the program in the attached black box.

### rRNA_sensor

The `rRNA_sensor` program compares input sequences to a `blastn` database of verified rRNA sequences using `blastn`. The program takes nine command-line arguments specified in Table 2. Each input sequence is classified into one of five classes based on its length and `blastn` results. A sequence is classified as *too long* or *too short* if its length is greater than the maximum length or less than the minimum length specified in the command by the user. To allow partial sequences and flexibility in the length, GenBank indexers were typically using a length interval of [400,2500] nt for prokaryotic 16S SSU rRNA. Empirical analysis shows that more than 99.8% of the full-length validated prokaryotic sequences have lengths in the range [900,1800], so this narrower range is recommended if one wants to check that sequences are typically full-length sequences.

**Table 2.** Command-line arguments for rRNA sensor

| argument index | argument name | description |
| --- | --- | --- |
| 1 | min_length | lower bound on sequence length |
| 2 | max_length | upper bound on sequence length |
| 3 | seq_file | input sequence file in FASTA format |
| 4 | output_file_name | name for summary output file |
| 5 | min_id_percentage | lower bound on percent identity |
| 6 | max_Evalue | upper bound on E-value |
| 7 | nprocessors | number of threads for blastn |
| 8 | output_dir | output directory path |
| 9 | blastdb | blastn database |

Sequences within the allowed length range are classified as either *no* if there are zero `blastn` hits, *yes* if they have at least one `blastn` hit that has an E-value of 1e-40 or less and a percent identity of 80% or more, or *imperfect_match* if there is at least one hit but the E-value or percent identity thresholds are not met for any hits. Sequences that are too long are probably either incorrect or containing extra flanking sequence that should be trimmed, while sequences that are too short may be valid partial sequences. The other tests based on quality of `blastn` matches codify the tests that GenBank indexers were doing internally before `rRNA_sensor` was implemented. Submitted sequences of a suitable length now classified as *no* would have been rejected in the past framework; sequences now classified as *yes* would have been accepted into GenBank in the past framework. In the current testing framework, `rRNA_sensor` is used as part of the `ribosensor` program as described below, not by itself.

There are two target `blastn` databases included with `rRNA_sensor`, one for prokaryotic 16S SSU rRNA and one for eukaryotic 18S SSU rRNA. The prokaryotic database includes 1267 sequences, 1205 of which are bacterial and the remaining 62 are archaeal. The eukaryotic database includes 1091 sequences. Additional, user-created `blastn` databases can also be used with the program. The prokaryotic database was updated most recently on June 29, 2017 by filtering and clustering the pre-existing database of 18,816 sequences used by GenBank indexers for 16S SSU rRNA analysis. One could repeat the same procedure with the larger version of the 16S SSU rRNA database described in Results. The initial database was filtered to remove 26 sequences outside the length range [900,1800]. The remaining 18,790 sequences were clustered using `UCLUST` [37] so that the surviving sequences were no more than 90% identical, leaving 1267 sequences. The eukaryotic 18S SSU rRNA database of 1091 sequences was updated most recently on September 27, 2018 by running version 0.28 of the Ribovore program `ribodbmaker` on an input set of 579,279 GenBank sequences returned from the eukaryotic SSU rRNA E-utilities (`eutils`) query provided in Results and discussion with command-line options --`skipfribo1` --`model SSU.Eukarya` --`ribo2hmm`.

### ribotyper

The `ribotyper` program is also designed to validate ribosomal RNA sequences but it differs from `rRNA_sensor` in the method of sequence comparison and the taxonomic breadth over which it applies. Instead of using `blastn, ribotyper` uses a profile HMM and optionally a covariance model (CM) to compare against input sequences. The profile HMM and CMs were built either from Rfam rRNA seed alignments (see Table 3) or from alignments created specifically for Ribovore by the authors for taxonomic groups not covered by the Rfam models.

**Table 3.** Profile models used by Ribovore. ’#seqs’ is the number of sequences in the multiple alignment used to build the model. ’length’ is the number of reference model positions. Abbreviations in ’taxonomy group’ column: ’Bac’ is Bacteria, ’Euk’ is Eukarya and ’Mito’ is Mitochondria.

| model name | gene | taxonomy group | #seqs | length | Rfam |
| --- | --- | --- | --- | --- | --- |
| SSU_rRNA_archaea | SSU rRNA | Archaea | 86 | 1477 | RF01959 |
| SSU_rRNA_bacteria | SSU rRNA | Bacteria | 99 | 1533 | RF00177 |
| SSU_rRNA_eukarya | SSU rRNA | Eukarya | 91 | 1851 | RF01960 |
| SSU_rRNA_microsporidia | SSU rRNA | Euk-Microsporidia | 46 | 1312 | RF02542 |
| LSU_rRNA_archaea | LSU rRNA | Archaea | 91 | 2990 | RF02540 |
| LSU_rRNA_bacteria | LSU rRNA | Bacteria | 102 | 2925 | RF02541 |
| LSU_rRNA_eukarya | LSU rRNA | Eukarya | 88 | 3401 | RF02543 |
| SSU_rRNA_mitochondria_metazoa | SSU rRNA | Mito-Metazoa | 83 | 954 | - |
| SSU_rRNA_mitochondria_amoeba | SSU rRNA | Mito-Amoeba | 2 | 1861 | - |
| SSU_rRNA_mitochondria_chlorophyta | SSU rRNA | Mito-Chlorophyta | 2 | 1200 | - |
| SSU_rRNA_mitochondria_fungi | SSU rRNA | Mito-Fungi | 4 | 1603 | - |
| SSU_rRNA_mitochondria_kinetoplast | SSU rRNA | Mito-Kinetoplast | 3 | 624 | - |
| SSU_rRNA_mitochondria_plant | SSU rRNA | Mito-Plant | 4 | 1951 | - |
| SSU_rRNA_mitochondria_protist | SSU rRNA | Mito-Protist | 2 | 1677 | - |
| SSU_rRNA_chloroplast | SSU rRNA | Chloroplast | 94 | 1488 | - |
| SSU_rRNA_chloroplast_pilostyles | SSU rRNA | Chloroplast | 1 | 1531 | - |
| SSU_rRNA_cyanobacteria | SSU rRNA | Bac-Cyanobacteria | 49 | 1487 | - |
| SSU_rRNA_apicoplast | SSU rRNA | Euk-Apicoplast | 3 | 1463 | - |

Sequence processing by `ribotyper` proceeds over two main stages. In stage 1, each sequence is compared against all profiles using a truncated version of the HMMER3 pipeline [38] optimized for speed. Only the first three stages of the HMMER3 pipeline are employed to compute a score for each sequence/profile comparison but without calculating accurate alignment endpoints. For each sequence, the best-scoring model is selected and used in the second stage where the HMMER3 pipeline is used again but this time in its entirety to compute likely endpoints of high-scoring hits to each model. These two stages are very similar to the classification and coverage determination stages of the VADR software package for viral sequence annotation [39]. The results of the stage 2 comparison are then post-processed to determine if any *unexpected features* exist for each sequence. There are 16 types of unexpected features, listed in Table 4.

**Table 4.**
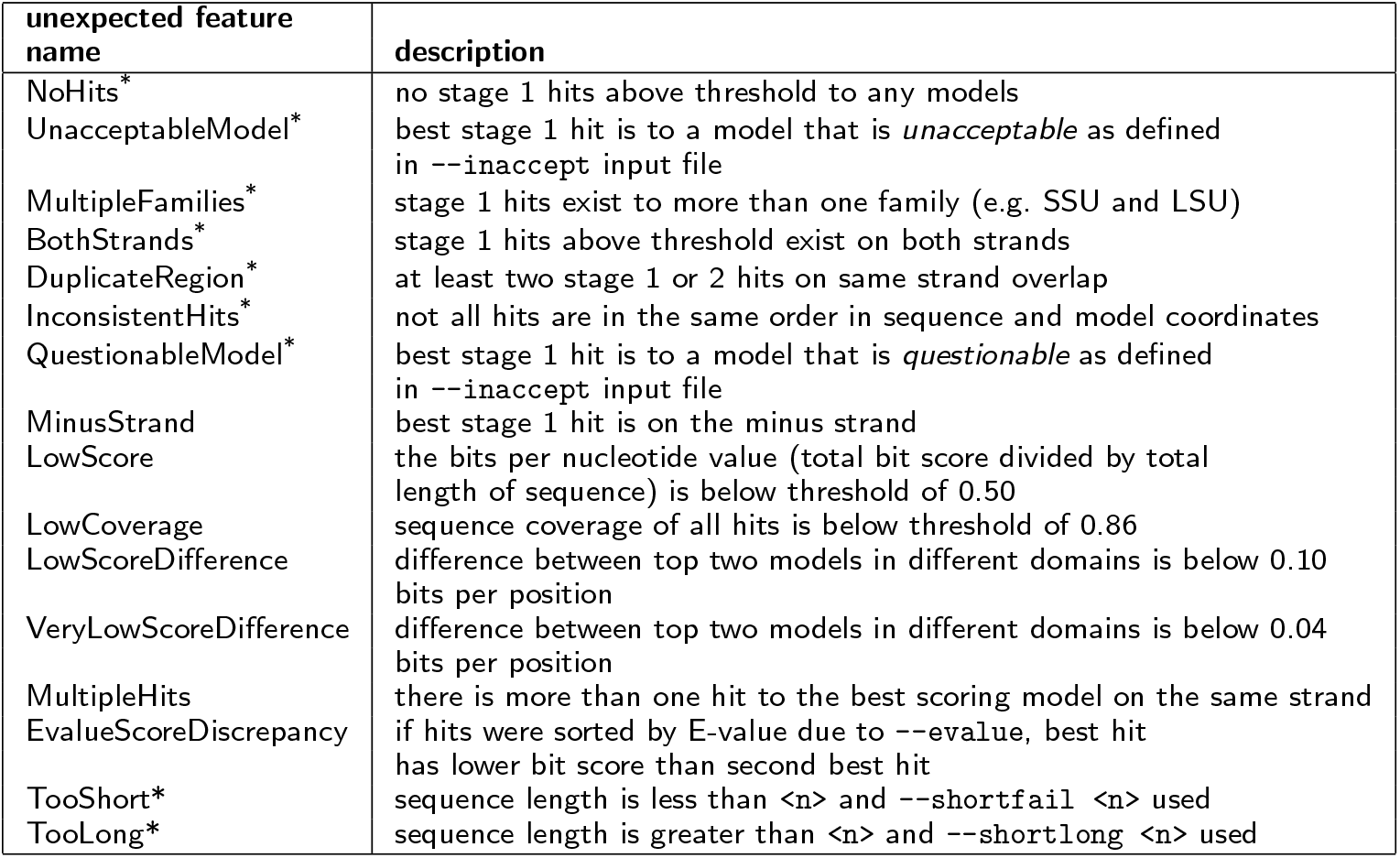
Attributes of the 16 types of ribotyper unexpected features. Unexpected features labelled with * in the first column are fatal by default, in that they cause a sequence to fail. UnacceptableModel and QuestionableModel can only potentially be reported if the --`inaccept` option is used. EvalueScoreDiscrepancy can only be reported if the --`evalues` option is used. TooShort and TooLong can only be reported if the --`shortfail` or --`longfail` options are used, respectively.

### ribosensor

The `ribosensor` program is a wrapper script that runs both `ribotyper` and `rRNA_sensor` and combines the results to determine if each sequence should pass or fail. This script was motivated partly by an effort to ease the transition for GenBank indexers between the pre-existing `blastn`-based system and a system based on profiles. Additionally, in some cases, the profile models in `ribotyper` allow some valid rRNA sequences that would fail `blastn` and `rRNA_sensor` to pass, and conversely some valid sequences pass `rRNA_sensor` and fail `ribotyper`, making a combination of the two programs potentially more accurate.

The ribosensor program can be run in one of two modes: 16S mode is the default mode and should be used for bacterial and archaeal 16S SSU rRNA sequences, and 18S mode should be used (by specifying the option -`m 18S` on the command-line) for eukaryotic 18S SSU rRNA. All sequences are first processed by ribotyper using command-line options --`scfail` --`covfail` --`tshortcov 0.80` --`tshortlen 350` to fail sequences for which LowScore and LowCoverage unexpected features are reported, and to specify that the threshold for LowCoverage is 80% for sequences of 350 nt or less. These options were selected based on results of internal testing by GenBank indexers. Next, `rRNA_sensor` is run, potentially up to three separate times, on partitions of the input sequence file separated based on length and using custom thresholds for each length range. Sequences that are shorter than 100 nt or longer than 2000 nt are considered too short or too long and are not analyzed. For sequences between 100 and 350 nt, a minimum percent identity of 75% and minimum coverage of 80% is enforced. For sequences between 351 and 600 nt, the minimum thresholds used are 80% percent identity and 86% coverage, and for sequences between 601 and 2000 nt the minimum thresholds used are 86% percent identity and 86% coverage. These thresholds can be changed via command-line options.

The results of `ribotyper` and `rRNA_sensor` are combined and each sequence is separated into one of four outcome classes depending on whether it passed or failed each program: *RPSP* (passed `ribotyper` and `rRNA_sensor`), *RPSF* (passed `ribotyper` and failed rRNA_sensor), *RFSP* (failed `ribotyper` and passed `rRNA_sensor`), and *RFSF* (failed both). Additionally, the reasons for failing each program are reported. For `ribotyper`, these are the unexpected features described above, each prefixed with a “R_” (e.g. R_MultipleFamilies). The possible errors for `rRNA_sensor` are listed in Table 5 and the possible errors for `ribotyper` are listed in Table 6. Finally, these errors are mapped to a different set of errors created for use within the pre-existing context of GenBank’s sequence processing pipeline shown which has its own error naming and usage conventions. This mapping is shown in Table 7. The “fails to” column is of practical importance because it indicates which errors cause a submission to not be accepted. If a submission fails to the submitter, then the submitter is expected to make corrections. In contrast, if a submission fails to indexer, then GenBank personnel review the error and determine if the submission can be corrected without further intervention by the submitter. If not, the submitter is contacted and asked for more information or to make the necessary corrections. More positively, if a submitter runs ribosensor before actually trying to submit and the submitter sees that the errors in the first seven rows and the third column of Table 7 do not occur, then, assuming the metadata for the submission are complete and valid, the submitter can have confidence that the submission to GenBank will be accepted.

**Table 5.**
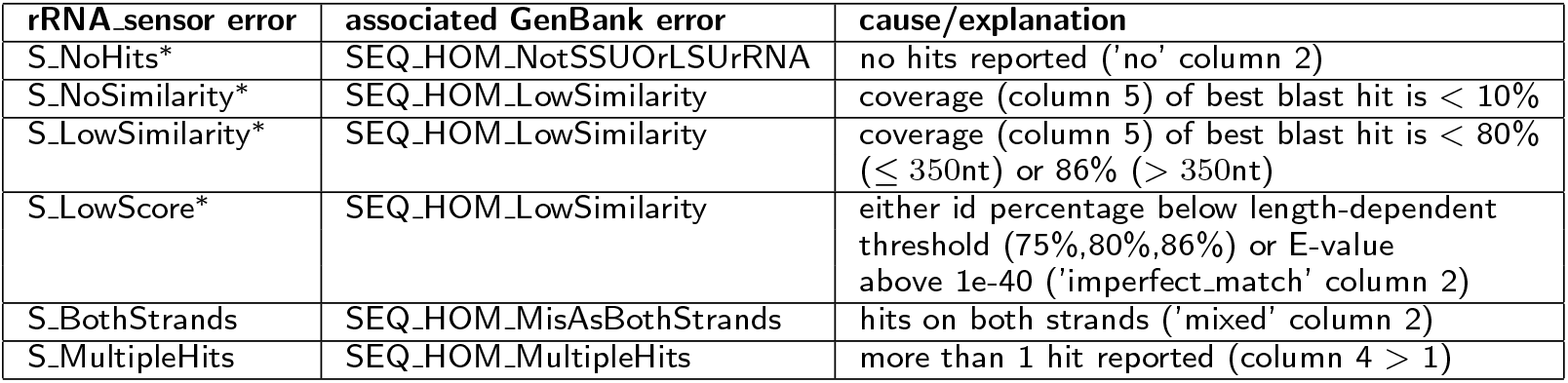
Descriptions of `rRNA_sensor` errors within `ribosensor` and mapping to the GenBank errors they trigger. ’*’: The first four `rRNA_sensor` errors do not trigger GenBank errors and are ignored by `ribosensor` if either (a) the sequence is ’RPSF’ (passes `ribotyper` and fails `rRNA_sensor`) and the -`c` option is *not* used with `ribosensor` or (b) the sequence is ’RFSF’ (fails both `ribotyper` and `rRNA_sensor`) and R_UnacceptableModel or R_QuestionableModel `ribotyper` errors are also reported.

**Table 6.** Descriptions of `ribotyper` errors within `ribosensor` and mapping to the GenBank errors they trigger. ’+’: these errors errors do not trigger a GenBank error if sequence is ’RFSP’ (fails `ribotyper` and passes `rRNA_sensor`);

| ribotyper error | associated GenBank error | cause/explanation |
| --- | --- | --- |
| R_NoHits | SEQ_HOM_NotSSUOrLSUrRNA | no hits reported |
| R_MultipleFamilies | SEQ_HOM_SSUAndLSUrRNA | SSU and LSU hits |
| R_LowScore | SEQ_HOM_LowSimilarity | bits/position score is < 0.5 |
| R_BothStrands | SEQ_HOM_MisAsBothStrands | hits on both strands |
| R_InconsistentHits | SEQ_HOM_MisAsHitOrder | hits are in different order in sequence and model |
| R_DuplicateRegion | SEQ_HOM_MisAsDupRegion | hits overlap by 10 or more model positions |
| R_UnacceptableModel | SEQ_HOM_TaxNotExpectedSSUrRNA | best hit is to model other than expected set<br>16S expected set: SSU.Archaea, SSU.Bacteria, SSU.Cyanobacteria, SSU.Chloroplast<br>18S expected set: SSU.Eukarya |
| R_LowCoverage | SEQ_HOM_LowCoverage | coverage of all hits is < 0.80 (if $\leq 350nt$) or 0.86 (if > 350nt) |
| R_QuestionableModel<sup>+</sup> | SEQ_HOM_TaxQuestionableSSUrRNA | best hit is to a 'questionable' model (if mode is 16S: SSU.Chloroplast) |
| R_MultipleHits<sup>+</sup> | SEQ_HOM_MultipleHits | more than 1 hit reported |

**Table 7.** Mapping of GenBank errors to the `rRNA_sensor` and `ribotyper` errors that trigger them. There are two classes of exceptions marked by two different superscripts in the table: ’*’: these `rRNA_sensor` errors do not trigger a GenBank error if: (a) the sequence is ’RPSF’ (passes `ribotyper` and fails `rRNA_sensor`) and the -`c` option is *not* used with `ribosensor`. or (b) the sequence is ’RFSF’ (fails both `ribotyper` and `rRNA_sensor`) and R_UnacceptableModel or R_QuestionableModel are also reported. ’+’: these `ribotyper` errors do not trigger a GenBank error if sequence is ’RFSP’ (fails `ribotyper` and passes `rRNA_sensor`);

| GenBank error | fails to | triggering rRNA_sensor/ribotyper errors |
| --- | --- | --- |
| SEQ_HOM_NotSSUOrLSUrRNA | submitter | S_NoHits*, R_NoHits |
| SEQ_HOM_LowSimilarity | submitter | S_NoSimilarity*, S_LowSimilarity*, S_LowScore*, R_LowScore |
| SEQ_HOM_SSUAndLSUrRNA | submitter | R_MultipleFamilies |
| SEQ_HOM_MisAsBothStrands | submitter | S_BothStrands, R_BothStrands |
| SEQ_HOM_MisAsHitOrder | submitter | R_InconsistentHits |
| SEQ_HOM_MisAsDupRegion | submitter | R_DuplicateRegion |
| SEQ_HOM_TaxNotExpectedSSUrRNA | submitter | R_UnacceptableModel |
| SEQ_HOM_TaxQuestionableSSUrRNA | indexer | R_QuestionableModel<sup>+</sup> |
| SEQ_HOM_LowCoverage | indexer | R_LowCoverage |
| SEQ_HOM_MultipleHits | indexer | S_MultipleHits, R_MultipleHits<sup>+</sup> |

### riboaligner

The `riboaligner` program was designed to help GenBank indexers to evaluate whether ribosomal RNA sequences are full length and do not extend past the boundaries of the gene. One application for a set of full length rRNAs is as part of the `blastn` database for screening and validating incoming sequences using `rRNA_sensor`, `ribosensor` or other `blastn`-based methods.

`riboaligner` first calls `ribotyper` to determine the best matching model for each sequence using special command-line options. The --`minusfail`, --`scfail` and --`covfail` options are used to specify that sequences with unexpected features of MinusStrand, LowScore and LowCoverage will fail. Additionally, the --`inaccept` *<f>* option is used to specify that the names of the desired models to use are in file *<f>*; only sequences that match best to one of these models is eligible to pass. The default set of acceptable models is SSU.archaea and SSU.bacteria by default. All sequences that score best to one of the acceptable models are aligned to that model using the `cmalign` program of Infernal which takes into account both sequence and secondary structure conservation. The alignment is then parsed to determine the length classification of each sequence based on the alignment. There are 13 possible length classes which are defined based on whether the alignment of each sequence extends to or past the first and final model reference position as well as how many insertions and deletions occur in the first and final ten model reference positions. More information on these classes can be found in the Ribovore documentation.

Only sequences that pass `ribotyper` will be aligned by `riboaligner`, and the per-sequence `ribotyper` pass/fail designation is not changed by `riboaligner`. The `riboaligner` summary output file is identical to the `ribotyper` output summary file with additional per-sequence information on the length class, start and stop model reference position of each aligned sequence and number of insertions/deletions in the first and final ten model positions.

### ribodbmaker

The `ribodbmaker` program is designed to create high quality datasets of rRNA sequences, which may be useful as reference datasets or `blastn` databases. It takes as input a set of candidate sequences and a specified rRNA model (e.g. SSU.Bacteria) and applies numerous quality control tests or filters such that only high quality sequences pass. The program performs the following steps:

1. fail sequences with too many ambiguous nucleotides (where “too many” is defined as more than the smaller of 5 and 0.5% the length of the sequence, but these thresholds can be changed by command-line options --`famaxn` and --`famaxf`)
2. fail sequences that do not have a specified species taxid in the NCBI taxonomy database
3. fail sequences that have non-weak `vecscreen` hits, suggesting the presence of vector contamination, as calculated by the VecScreen plus taxonomy software package [40]
4. fail sequences that have unexpected internal repeats as determined by comparing each sequence against itself using `blastn` and finding off-diagonal local alignments with an E-value of no more than 1 and length at least 20 for the plus strand and 50 for the minus strand
5. fail sequences that fail `ribotyper`, including matching best to a model other than the specified one, using non-default options --`minusfail` --`lowppossc 0.5` --`scfail` to specify that sequences with best hits on the minus strand or with scores below 0.5 bits per nucleotide will fail
6. fail sequences that fail `riboaligner`, including matching best to a model other than the specified one, using non-default options --`lowppossc 0.5` --`tcov 0.99` to specify that sequences with scores below 0.5 bits per nucleotide or for which less than 99% of the sequence length is covered by hits will fail
7. fail sequences that do not cover a specified span of model positions (are too short)
8. fail sequences that survive all above steps but do not meet expected criteria of an *ingroup* analysis based on taxonomy and alignment identity

In step 6, `riboaligner` outputs multiple sequence alignments of all sequences. These alignments are used for further scrutiny of each sequence in step 7, the *ingroup* analysis step. At this stage, sequences that do not cluster (based on alignment identity) with other sequences in their taxonomic group fail. Finally, sequences that survive all stages are clustered based on alignment identity and centroids for each cluster are selected for the final set of surviving sequences.

Steps 2, 3, and 8 require access to the NCBI taxonomy database and further that each input sequence be assigned in the nucleotide database to a unique organism in the taxonomy database. This restricts the use of `ribodbmaker` to sequences already present in GenBank. The taxonomy criterion excludes, for example, some chimeric sequences that have been engineered and patented. Users can run ribodbmaker on other sequences, but must bypass these steps using the --`skipftaxid`, --`skipfvecsc`, and --`skipingrup`. The VecScreen plus taxonomy package is only available for Linux and so is not installed with Ribovore on Mac/OSX. Consequently, the following `ribodbmaker` options must be used on Mac/OSX: --`skipftaxid` --`skipfvecsc` --`skipingrup` --`skipmstbl`. In general, `ribodbmaker` is highly customizable via command-line option usage, and can be run using many different subsets of tests. For more information on command-line options see the `ribodbmaker`.md file in the Ribovore documentation subdirectory.

As described above, `riboaligner` calls `ribotyper`, so `ribotyper` is actually called twice by `ribodbmaker`, once in step 5 and once in step 6. In the `riboaligner` step, `ribotyper` is called with options that differentiate its usage from step 5, making the criteria for passing more strict in several ways. The --`difffail` and --`multfail` options are used to specify that sequences with unexpected features of LowScoreDifference and VeryLowScoreDifference will fail. Additionally, a CM is used instead of a profile HMM for the second stage (`ribotyper` --`2slow option`) and any sequence for which less than 99% of the nucleotides are covered by a hit in the second stage will fail (--`tcov 0.99` option). Finally, the --`scfail` option, which is used in the `ribotyper` call in step 5, is not used in step 6.

### Ribovore reference model library and blastn databases

The Ribovore package includes 18 sequence- and structure-based alignments and corresponding CMs, listed in Table 3. The alignment and CM files are included when Ribovore is installed and are also available on GitHub. Seven of the 18 alignments are from Rfam, and the other 11 were created during development of the package. `rRNA_sensor` includes two `blastn` databases: one of 1267 bacterial and archaeal 16S SSU rRNA sequences created by clustering and filtering the `blastn` database already in use at GenBank in 2017 when development of the script began, and one of 1091 eukaryotic 18S SSU rRNA sequences created by filtering a sequence dataset generated by `ribodbmaker`.

All 18 of the Ribovore model alignments are the end products of a multi-step model refinement procedure using the valuable secondary structure data available from CRW [22] and sequences from GenBank. For each gene and taxonomic group (e.g. SSU rRNA eukarya), an initial alignment with consensus secondary structure was created based on combining alignments and individual sequence secondary structure predictions from CRW as described in [20, 41], and used to build a CM using the Infernal program `cmbuild`. That CM was then calibrated for database sequence search using `cmcalibrate` and searched against all currently available rRNA sequences in GenBank. The resulting high-scoring hits were then filtered for redundancy and manually examined and surviving sequences were realigned to the model to create a new alignment. In some cases the consensus secondary structure was modified slightly based on the new alignment. Some models were further refined by additional iterations of building, searching, and realigning.

Eight of the 18 Ribovore models are SSU models with fewer than 10 sequences in the training alignment (Table 3). These are for taxonomic groups with relatively few known example sequences for which the consensus secondary structure is distinct but not as well understood as for other groups, like 16S SSU rRNA. Six of these eight are non-metazoan mitochondrial models, one is a chloroplast model for the *Pilostyles* plant genus, and one is for apicoplasts. These eight models are less mature than the other ten models, but they are included in the package for completeness and we plan to improve them in future versions. Currently, users should be cautious when interpreting results that involve any of these eight models.

From each of the 18 Ribovore model alignments, two separate CMs were constructed using different command-line parameters to the `cmbuild` program of Infernal. One model was built using `cmbuild`’s default entropy weighting feature that controls the average entropy per model position [14, 20], and one was built using the `cmbuild --enone` option, which turns off entropy weighting. The non-entropy weighted models, which perform better at sequence classification in our internal testing (results not shown), are used by `ribotyper`, and the entropy weighted models are used by `riboaligner` for sequence alignment because they are slightly more accurate at getting alignment endpoints correct based on our own internal test results (not shown).

### Timing measurements

Timing measurements of `rRNA_sensor`, `ribotyper`, `riboaligner`, and `ribodbmaker` were done primarily on an Intel(R) Xeon(R) Gold 5118 CPU @ 2.30GHz with 48 cores and running the CentOS7.8.2003 version of Linux. We used one thread except for tests of `rRNA_sensor` that measured the effect on wall-clock time of increasing the number of threads. For the runs of `ribodbmaker`, we used the NCBI compute farm to parallelize some time intensive steps. The reason for using the compute farm only for the `ribodbmaker` tests is that `ribodbmaker` is intended primarily for curation of databases at NCBI, while the other modules are intended to be used both by submitters around the world and GenBank indexers at NCBI.

## Results and discussion

Ribovore is used directly or indirectly by NCBI and GenBank in various ways: as part of its submission pipelines for rRNA sequences, through the BLAST Web server (https://blast.ncbi.nlm.nih.gov/Blast.cgi?PROGRAM=blastnPAGE_TYPE=BlastSearchLINK_LOC=blasthome) and by facilitating the validation of sequences from type material to be incorporated into new records in the RefSeq database (https://www.ncbi.nlm.nih.gov/bioproject/224725). We detail each of these uses below and then compare the capability of Ribovore for fungal rRNA sequence validation to related projects.

### rRNA sequence submission checking

Submitters of rRNA sequences to GenBank who use the NCBI Submission Portal can choose between 12 different subtypes, listed in Table 8. For most submission subtypes, the sequences are analyzed via a `blastn`-based pipeline by comparing each submitted sequence against a `blastn` database for the specific submission subtype. Three of these submission subtypes (ITS1 and ITS2 and 16S-23S IGS) are for non-rRNA sequences. For four of the remaining nine subtypes, the `blastn` database currently used was created with the help of `ribodbmaker`, as discussed more below. The `ribosensor` program is used instead of the `blastn` pipeline to analyze 16S prokaryotic SSU rRNA submissions of 2500 or more sequences for which the submitter chooses the attribute *uncultured* to describe the sequences.

**Table 8.** NCBI rRNA and ITS sequence submission types and attributes. The ’submission type’ column indicates the three possible rRNA sequence related options for GenBank submissions available at https://submit.ncbi.nlm.nih.gov/subs/genbank/. There is not an intergenic spacer type of eukaryotes at this time. The ’submission subtype’ column indicates the more specific possible sequence types a submitter can select after choosing one of the options in the first column. The ’validation method’ column indicates whether incoming sequences are processed with ribosensor, blastn using a database constructed with ribodbmaker (’blast(ribo)’) or blastn using either the general non-redundant (nr) database or a database constructed by other means (’blastn’). The ’percentage of accepted submissions’ and ’percentage of accepted sequences’ reflect rRNA/IGS/ITS submissions published between Jan 1, 2020 and May 31, 2020. Note that the percentages for rows 3 and 4 are summed and reported in row 3, and for rows 5 through 12 (all eukaryotic submission types) are summed and reported in in row 5. Counts pertain only to submissions that advanced through enough preliminary checks to be assigned an internal submission code.

| submission type | submission subtype | validation method | percentage of accepted submissions | percentage of accepted sequences |
| --- | --- | --- | --- | --- |
| Prokaryotic rRNA/IGS | SSU rRNA only (16S) ≥ 2500 seqs | ribosensor | 0.648% | 99.604% |
| | SSU rRNA only (16S) $<$ 2500 seqs | blastn(ribo) | 43.934% | 0.1252% |
|  | LSU rRNA only (23S) | blastn(ribo) | 0.754% | 0.0027% |
|  | intergenic spacer (16S-23S IGS) | blastn | (sum of 2 rows) |  |
| Eukaryotic Nuclear rRNA/ITS | contains rRNA-ITS region | blastn | 54.663% | 0.2682% |
|  | SSU rRNA only (18S) | blastn(ribo) | (sum of 8 eukaryotic rows) |  |
|  | LSU rRNA only (28S) | blastn(ribo) |  |  |
|  | ITS1 only | blastn |  |  |
|  | ITS2 only | blastn |  |  |
| Eukaryotic Organellar rRNA | mitochondrial SSU rRNA (12S) | blastn |  |  |
|  | mitochondrial LSU rRNA (16S) | blastn |  |  |
|  | chloroplast SSU rRNA (16S) | blastn |  |  |
|  | chloroplast LSU rRNA (23S) | blastn |  |  |

For `ribosensor`, the default parameters are used to determine if sequences should pass or fail as discussed in the Implementation section. For `blastn`, sequences are evaluated based on the average percentage identity, the average percentage query coverage, and the percentage of gaps in the alignments for the top target sequences. Additionally, using a `blastn`-based method that predates and inspired `rRNA_sensor`, sequences that are suspected to be misassembled or incorrectly labelled taxonomically fail. Specifically, the query sequence is tested with `blastn` against the 16S SSU rRNA database described below and the matches are ranked in increasing order of E-value. A sequence passes the misassembly test if and only if the best matches each have exactly one local alignment. The taxonomy tests are based on a comparison of the proposed taxonomy from the submitter and the taxonomic information of the top matches, taking into account variant spellings and synonyms in NCBI Taxonomy. The exact thresholds for these pre-Ribovore, `blastn`-related comparisons vary according to the type of submission and are outside the scope of this paper.

For both the `blastn` and `ribosensor` pipelines submissions in which all sequences pass and which have the required metadata are automatically deposited into GenBank. All other submissions fail and are either sent back to the submitter with automated error reports or manually examined further by GenBank indexers, depending on the specific reason for the failure.

A key objective of distributing the Ribovore software is to permit submitters to do on their own computers similar checks to those done by the GenBank submission pipeline. In 2018, we began using earlier versions of `ribosensor` to analyze large-scale 16S prokaryotic SSU rRNA submissions. This remains the only submission type for which `ribosensor` is employed in an automated way, although we plan to expand to additional genes and taxonomic domains in the future. Parts of Ribovore are also used manually by GenBank indexers to evaluate some submissions.

The most common type of rRNA submission by far is 16S prokaryotic SSU rRNA (Table 8). Between July 1, 2018 and May 31, 2020, 33,388 submissions of 16S SSU rRNAs with less than 2500 sequences (or for which the submitter indicated the sequences were from cultured organisms), were handled by the `blastn` pipeline. The total number of sequences in these submissions was 240,112, for an average of 7.2 sequences per submission. In the same time interval, `ribosensor` processed 242 16S SSU rRNA submissions comprising 49,868,017 sequences for an average of 206,066.2 sequences per submission. In the first six months of 2020, `ribosensor` processed more than 99.6% of the sequences deposited in GenBank via any of the the rRNA or ITS submission pipelines (Table 8).

### Construction and usage of rRNA databases for blastn

Four of the ten rRNA `blastn` databases used for submission checking were created by `ribodbmaker` as indicated in Table 8. An additional three `blastn` databases, available in the web server version of BLAST are described below. All `blastn` databases we mention can be retrieved for local use from the directory https://ftp.ncbi.nlm.nih.gov/blast/db/ The databases are (re)generated semi-automatically by extracting large sets of plausible sequences from Entrez and providing them as input to `ribodbmaker`. The `ribodbmaker` program is run so that the tests for ambiguous nucleotides, specified species, vector contamination, self-repeats, `ribotyper`, `riboaligner`, and model span are all executed. However, the databases are allowed to contain more than one sequence per taxid and the ingroup analysis is skipped (--`skipingrup` option). Only sequences that pass `ribodbmaker` tests are eligible to be in the `blastn` databases. To keep the 16S prokaryotic SSU, eukaryotic SSU, and eukaryotic LSU databases to a reasonable size, the sequences that pass `ribodbmaker` are clustered with `UCLUST` [37] at a threshold of 97% identity and all other parameters at default values. The clustering stage of `ribodbmaker` is not used for this purpose and is skipped by using the --`skipclustr` option.

The 16S prokaryotic SSU rRNA BLAST database is generated starting from all sequences in the GenBank nucleotide database that match NCBI BioProject IDs PRJNA33175 or PRJNA33317 using the eutils query in the first row of Table 9. PRJNA33175 has the title “Bacterial 16S Ribosomal RNA RefSeq Targeted Loci Project”. PRJNA33317 has the title “Archaeal 16S Ribosomal RNA RefSeq Targeted Loci Project”. This formal query is supplemented by manual searches of the journal *International Journal of Systematics and Evolutionary Biology* (https://www.microbiologyresearch.org/content/journal/ijsem), where many new bacterial species are announced and peer-reviewed, along with their 16S SSU rRNA sequences. Among the databases described here, the 16S SSU rRNA database is the only one restricted to sequences from “type material” that have been more stringently vetted before curation for RefSeq. The fungal RefSeq records described in a later subsection are also restricted to be from “type material”.

**Table 9.**
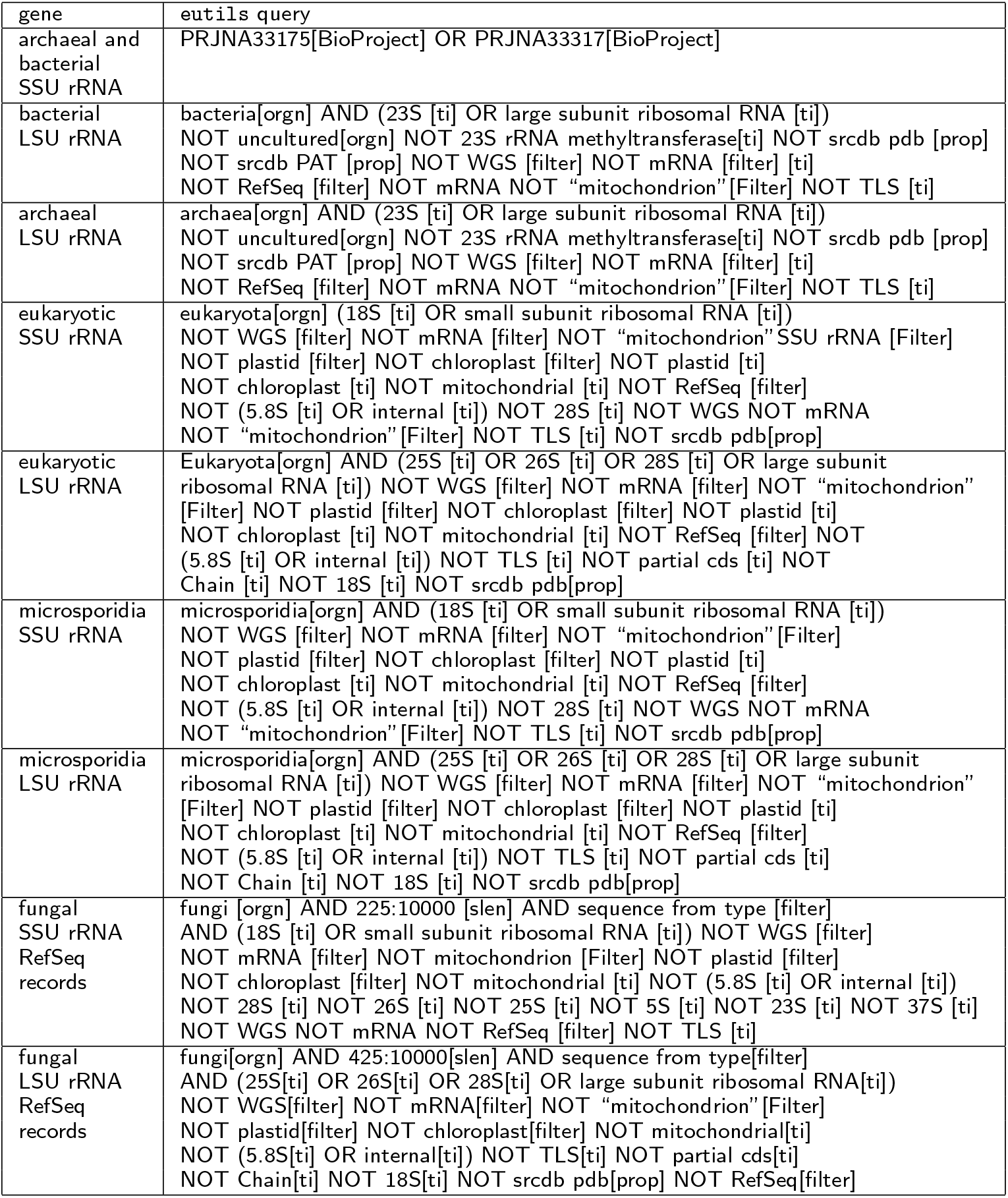
Queries used in command-line `eutils` to collect input datasets for `ribodbmaker`.

Table 9 also lists the eutils queries to the nucleotide database that are used for 23S prokaryotic LSU rRNA, eukaryotic SSU rRNA, eukaryotic LSU rRNA, *Microsporidia* SSU rRNA, and *Microsporidia* LSU rRNA. When we seek sequences that are likely to be complete, not larger genome pieces, and not partial, we add a constraint on the length with an extra term such as 1500:18000[slen] for eukaryotic SSU rRNA. The main attribute that distinguishes *Microsporidia* is that the lower bound on slen for complete sequences is set about 300-500 nucleotides lower as explained in Background. To find possibly partial LSU sequences that are long enough to cover the variable regions, we add the condition 425:1000[slen]. These queries rely on standardized nomenclature and structure of the *definition line* of GenBank sequence records which contain information about the source organism, feature content, completeness and location. Since 2016, these definition lines have been constructed formulaically during the processing of submissions. For example, the sequence MT981756.1 has the title: “Staphylococcus epidermidis strain RA13 16S ribosomal RNA gene, partial sequence”, and the sequence MN158348.1 has the title “Tetrahymena rostrata strain TRAUS 18S ribosomal RNA gene, internal transcribed spacer 1, 5.8S ribosomal RNA gene, internal transcribed spacer 2, and 28S ribosomal RNA gene, complete sequence”.

### Creation of fungal rRNA RefSeq entries using ribodbmaker

NCBI’s RefSeq project seeks to create a representative, non-redundant set of annotated genomes, transcripts, proteins and nucleotide records including rRNA sequences [36]. Since 2018, `ribodbmaker` has been used to screen the set of fungal 18S SSU rRNA and 26S LSU rRNA sequences. Table 9 lists the queries used to identify candidates to be new fungal SSU and LSU rRNA RefSeq records.

Studies that target fungal rRNAs frequently attempt to obtain SSU rRNA sequences that span most of the V4 and part of the V5 variable regions, or LSU rRNA sequences that span the D1 and D2 variable regions as these have been shown to be phylogenetically informative [42, 43]. These regions correspond to Rfam RF01960 model positions 604 to 1070 (SSU) and RF02543 model positions 124 to 627 (LSU). Correspondingly, `ribodbmaker` is run with command-line options (--fmlpos and --fmrpos) that enforce that only sequences that span these model coordinates can pass. The following `ribodbmaker` options are used for SSU: --`fione` --`fmnogap` --`fmlpos 604` --`fmrpos 1070` -`f` --`model SSU.Eukarya` --`skipclustr`, and for LSU: --`fione` --`fmnogap` --`fmlpos 124` --`fmrpos 627` -`f` --`model LSU.Eukarya` --`skipclustr`.

### NCBI BLAST webpage rRNA target databases

For many years, NCBI has been offering searches of nucleotide and protein databases with various modules of BLAST [44] through the NCBI BLAST webpage. Most commonly, searches of nucleotide queries use a comprehensive “nonredundant (nr)” database of nucleotide sequences or databases of whole genomes. A disproportionate number of queries are rRNA sequences. When `blastn` users know that their queries are of these special types, searching smaller targeted databases that exclude sequences unexpected to have a significant match to the query reduces running time and leads to more focused results. The BLAST webpage now allows users to select from three `ribodbmaker`-derived rRNA target databases, listed in Table 10. The 16S SSU rRNA database is identical to the one used by the `blastn` submission pipeline. The other two are specific to *Fungi*, due to the popularity of the analysis of rRNA sequences for studies of that kingdom. The fungal SSU and fungal LSU BLAST databases are effectively equivalent to the sets of curated RefSeq records described below. The availability of these databases was announced in late 2019, and the number of `blastn` runs and unique `blastn` visitors who selected each database during the seven-month period November 12, 2019 - June 22, 2020 are reported in Table 10. The usage suggests that there is sufficient user demand to justify the curation effort. As of November 24, 2020, there are 2,836 fungal SSU RefSeq records and 7,573 fungal LSU RefSeq records, almost all of which were curated with Ribovore.

**Table 10.** Web `blastn` usage of specialized rRNA databases that are curated using Ribovore to decide which sequences are valid. Usage was measured during November 12, 2019 - June 22, 2020.

| database | runs | runs/day | visits | visits/day |
| --- | --- | --- | --- | --- |
| 16S rRNA sequences (SSU) from Bacteria and Archaea | 355,479 | 1921.5 | 92,638 | 500.7 |
| 18S rRNA sequences (SSU) from Fungi and reference type material | 5,724 | 30.9 | 2,876 | 15.5 |
| 28S rRNA sequences (LSU) from Fungi and reference type material | 4,535 | 24.5 | 1,861 | 10.1 |

### Comparison to curated sets of fungal rRNA sequences from Silva

We compared iteratively our `ribodbmaker` approach to curating fungal RefSeq records with other curatorial efforts that are part of the Silva project [27]. The purposes of this comparison were:

1. to identify new candidate RefSeq records and possible weakness in our procedures for choosing RefSeq records,
2. to test whether Ribovore works on curated data sets and to correct errors associated with some sequences in NCBI databases, such as misleading definition lines,
3. to characterize what proportion of sequences curated by others pass the Ribovore criteria and why sequences fail.

As noted above, fungal LSU sequences submitted since 2016 or 2017 should have definition lines that match this query. In the course of doing the SilvaParc LSU tests described below, we corrected the definition lines of 132 older sequences that are fungal LSU and passed all `ribodbmaker` tests, but did not match the above eutils query. The current fungal RefSeq SSU and LSU sequences can be obtained with the queries “PRJNA39195[BioProject]” and “PRJNA51803[BioProject]”, respectively or both from the ftp site: https://ftp.ncbi.nlm.nih.gov/refseq/TargetedLoci/Fungi/.

For our comparison of fungal sequences, we used a curated set of 8,770 SSU sequences from Silva [45], a set of 1,461 SSU sequences from Silva in the phylum *Microsporidia*, a set of 2,993 high-quality LSU reference sequences from Silva, and a much larger set of 394,247 sequences from Silva called Parc [26, 46]. We denote these four sets as *Yarza*, *SilvaMicrosporidia*, *SilvaRef*, and *SilvaParc*, respectively. To set up the SilvaMicrosporidia set, we downloaded the FASTA files for all of SilvaParc SSU and extracted 1,468 sequences labeled as being from the phylum *Microsporidia*; of these, 1,461 sequences were in GenBank with sufficient taxonomy information to be considered for `ribodbmaker`. SilvaParc contains fewer than 50 *Microsporidia* LSU sequences, supporting our previous assertion that the SSU has been much more studied than the LSU in *Microsporidia*.

Similarly, to set up the SilvaRef and SilvaParc sets, we downloaded FASTA files for all SilvaRef LSU sequences for all taxa in version 132 on July 23, 2020 and all LSU Parc sequences in version 132 on August 1, 2020. We filtered for all sequences that had the token “Fungi” in the definition line. A small number of sequences had to be dropped subsequently because 1) they are not from the kingdom *Fungi* (e.g., they may be from a pathogen of a fungus) 2) they were absent from the nuccore database of GenBank either due to being “unverified” or from certain types of patents or 3) due to phylogenetic discrepancies (cf. [47]) that we subsequently fixed as part of the first objective listed above. In all data sets, we retrieved the most recent version of all GenBank accessions, which differs from the curated version for a very small number of sequences since version 132 of Silva is recent. In our analysis of the SilvaParc set, 96 non-fungal sequences were inadvertently included in the analysis, and excluded only while checking the results. The results of the three comparisons are shown in Table 11.

**Table 11.**
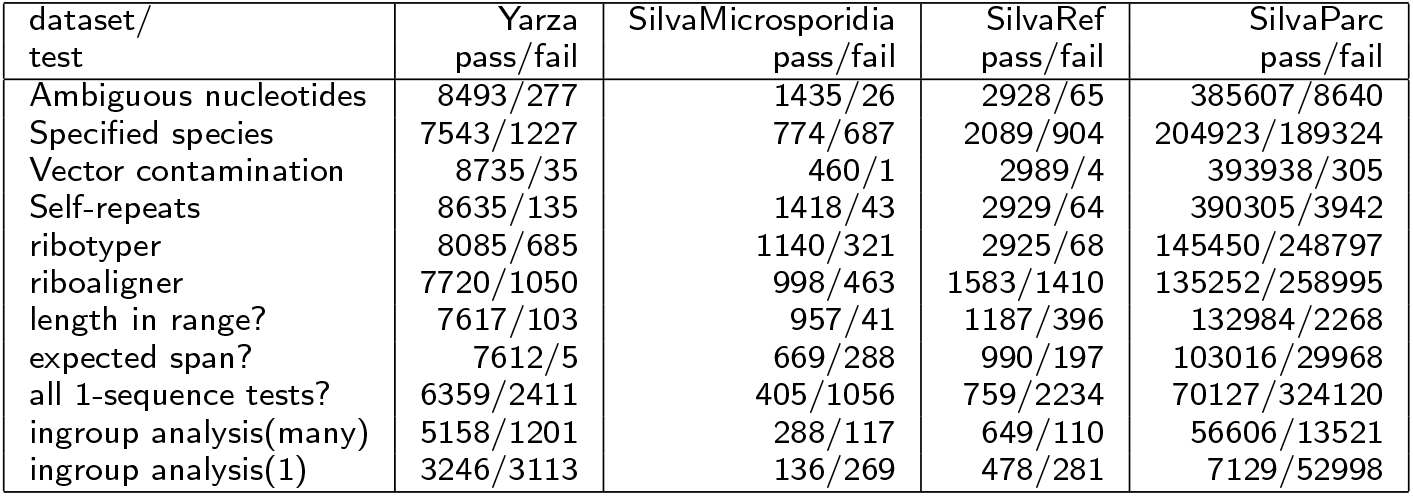
Summary of `ribodbmaker` pass/fail outcomes for Yarza(SSU), SilvaMicrosporidia(SSU), SilvaRef(LSU), and SilvaParc(LSU) datasets. All tests except the ingroup analysis depend only on the sequence being tested. The four tests for ambiguous nucleotides, specified species, vector contamination, and self-repeats are done on all sequences, so sequences may fail more than one test. Only sequences that pass the `ribotyper` test are eligible as input to `riboaligner`. Only sequences that pass the `riboaligner` test are eligible to be tested for length and alignment span. Only sequences that pass all 1-sequence tests are eligle for ingroup analysis. The ingroup analysis can be done allowing many sequences from the same taxon to pass or limiting to 1 the number of sequences that pass from each taxon (argument --`fione`). The many option is a more meaningful test; we show the 1 option just for comparison. Data and instructions for reproducing these comparisons are available at https://github.com/nawrockie/ribovore-paper-2021-supplementary-material.

The main steps in these tests consisted of:

1. Download and uncompress a FASTA file of source sequences from the supplementary information of [45] or from the Silva FTP retrieval site, which we denote `File1.fa`.
2. As explained in the Ribovore documentation, retrieve and condense the current version of NCBI’s taxonomy tree. An important and subtle column is the boolean (0/1) specified species column for each taxon; a 1 in this column for the row of taxon *t* means that according to NCBI’s Taxonomy Group the taxon name is valid and currently peferred; a 1 in this column is a necessary condition for a sequence from taxon *t* to be eligible to be in the rRNA databases or to be a RefSeq. Call the resulting file `taxonomy.txt`.
3. (For tests of Silva data only) Extract the definition lines for sequence identifiers of interest with the command: `grep Fungi File1.fa | grep -v Bacteria` or `grep Microsporidia File1.fa | grep -v Bacteria`, redirecting the output to an intermediate file. The second command in the pipe removes most sequences from fungal pathogens that are not actually from the kingdom of Fungi. For the Fungal SSU set, all sequences are of interest, so the simpler command `grep ‘‘>’’ File1.fa` extracts the definition lines.
4. Extract GenBank accessions without the versions from the definition lines at step 3. Versions are removed because for some sequences, the version in Silva has been superseded in GenBank with a newer version.
5. Use the NCBI package `eutils` [48] to retrieve from the nucleotide database of GenBank all currently live accessions from the accession sets derived at the previous step. Some sequences get dropped at this step because they are no longer live. Call the FASTA file at this step `File2.fa`.
6. Use the NCBI standalone tool srcchk (available at: ftp://ftp.ncbi.nih.gov/toolbox/ncbi_tools/converters/by_program/srcchk/) to check which sequences in `File2.fa` have a valid and fully consistent taxonomy entry. Remove sequences that do not get a normal result from `srcchk` because they will cause Ribovore to halt. A small number (well below 1%) of sequences get removed at this step either because they are engineered sequences from patents or because there are transient inconsistencies between the NCBI taxonomy tree and the organism values in the GenBank nucleotide records. Call the resulting file `File3.fa`.
7. Run `ribodbmaker --taxin taxonomy.txt --skipclustr --model <model-value> --fmlpos <left-boundary> --fmrpos <right-boundary> --fmnogap --fione --pidmax 71000 --indiffseqtax -f -p <output directory>`. The value of `<model-value>` was either SSU.Eukarya, SSU.Microsporidia, or LSU.Eukarya, depending on the test being done. The values of `<right-boundary>` and `<left-boundary>` are set in a model-specific manner according to the recommended values in the Ribovore documentation (`ribodbmaker`.md file in documentation subdirectory). The 71000 is an upper bound on the number of sequences input in our various tests. Version 0.40 of Ribovore was used.

For the large test of SilvaParc, we ran the last `ribodbmaker` step separately on various subsets of `File3.fa`, then collected all sequences that passed all sequence-specific tests, and did a final run of `ribodbmaker` to include the ingroup analysis. This split into multiple runs achieves better parallelism and throughput for large versions of `File3.fa` because the ingroup analysis is the only step in which the results for any single sequence depend on which other sequences are included in the input. The numbers of eligible sequences reported in Table 11 are those included in File3.fa. The results are slightly sensitive to changes in the NCBI taxonomy tree, which is updated daily. For the Yarza tests, we used the NCBI taxonomy tree as of August 1, 2020 and for the Silva tests we used the tree as of September 22, 2020.

It was not our intent to compare sets of “passing” sequences because the criteria for fungal RefSeq records are deliberately more stringent than for inclusion in Silva. Most notable are the two taxonomic tests: 1) that each sequence should come from a specified species and 2) that in selecting sequences for RefSeq, we may choose to keep only one sequence per species taxid, as specified by the command-line option --`fione`, to avoid redundancy.

Indeed, the analysis of fungal sequences from Silva yielded some new fungal RefSeq records; specifically, we added 4 SSU sequences from the Yarza set, and 10 LSU sequences from the SilvaRef and SilvaParc data sets. However, the tests results also show some possible improvements in the Silva curation. It appears that a small number of Silva sequences have vector contamination and more than 1% may be misassembled as indicated by self-repeats, which are not expected in fungal SSU and LSU rRNA genes (see Methods, subsection `ribodbmaker` for the self-repeat criteria). It appears that the Yarza SSU data set was carefully curated for sequences to be full-length and not too long, but in the SilvaRef and SilvaParc data sets more than 20% of sequences have either a length that is out of the range of typical eukaryotic LSU sequences or do not span the range [124,627] that includes the D1/D2 regions typically covered for species differentitation. Thus, it appears that the Silva resource curation could arguably be improved by checking sequence ends, so as to trim long sequences, remove short sequences, and remove sequences that are unlikely to be full LSU sequences. The sequences could be too long either because they were not trimmed to the LSU boundary or possibly because they contain introns. In the SilvaMicrosporidia test, we tested for the presence of the most conserved V4 and V5 regions only with a permissive expected span of [380,800].

### Partial comparison to RNAmmer

To our knowledge, there is no other software that solves the rRNA validation problem as we have formulated it for GenBank submissions. One widely used software package that solves a related problem is `RNAmmer` [28]. The problem that RNAmmer solves is to find likely rRNA sequences within larger sequences by using an old version of HMMER (v2.3.2) to compare against one of six profile HMMs. To accelerate searches, RNAmmer first utilizes a small *spotter* profile HMM that models only the most conserved 75 consecutive positions of the overall rRNA alignment to detect rRNA regions, padding those regions with extra sequence on each end, and then using a profile of the full rRNA to determine gene boundaries within those padded regions. So far as we could determine, RNAmmer does not work with the up-to-date HMMER version 3 nor with arbitratry profile HMMs or CMs.

Nevertheless, one can provide as input to RNAmmer FASTA files of putative rRNA sequences, pretending that they were larger contigs. The module of Ribovore that is closest in purpose to this usage of RNAmmer is `ribotyper`. To allow comparison between RNAmmer and `ribotyper`, one can define that a sequence *passes* RNAmmer if RNAmmer produces in the output at least one HMMER-based prediction for that sequence using the intended rRNA model (e.g., eukaryotic SSU for the Yarza set) and *fails* if there are zero such predictions. This comparison is unfair to RNAmmer because it does not use the predicted intervals, which are the most useful part of the output when RNAmmer is used with large contigs as inputs.

We compared the performance of `ribotyper` versus RNAmmer on the Yarza SSU set and the SilvaRef LSU set. We used the `ribotyper` results obtained from the `ribodbmaker` tests described above and summarized in Table 11. Among the 8,770 SSU sequences in the Yarza set: 7,999 passed both `ribotyper` and RNAmmer, 23 failed both `ribotyper` and RNAmmer, 665 passed RNAmmer and failed `ribotyper`, and 86 passed `ribotyper` and failed RNAmmer. Among the set of 665, 557 sequences include “internal transcribed spacer” or “ITS” in the definition line, and 69 of the other 108 have lengths above 5,000 nt, indicating that all but at least 39 of the sequences likely include sequence outside the SSU rRNA sequence (which is rarely more than 3Kb) and so are expected to fail `ribotyper`. Of the 86 sequences that passed `ribotyper` and failed RNAmmer, 46 of them would pass RNAmmer if the E-value and bit score thresholds for the spotter HMM which are hard-coded at 1E-5 and 0 were changed to 1 and −100 in the rnammer Perl script, indicating that these sequences do not match well to the spotter profile HMM used for eukaryotic SSU rRNA.

Among the 2,993 LSU sequences in the SilvaRef set: 2,481 passed both RNAmmer and `ribotyper`, 21 failed both RNAmmer and `ribotyper`, 47 passed RNAmmer and failed `ribotyper`, and 444 passed `ribotyper` and failed RNAmmer. Among the 47 sequences that passed RNAmmer and failed `ribotyper`, 44/47 can be explained because they have one of three errors that `ribotyper` looks for and would not necessarily lead RNAmmer to have no output matches: R_DuplicateRegion (24 sequences), R_BothStrands (7 sequences), R_MultipleFamilies (13 sequences). Many of the 47 sequences are described on the definition lines as a “shotgun assembly”; accordingly, the R_DuplicateRegion and R_BothStrands errors indicate two different errors that occur commonly in assembling nucleotide sequences into contigs. Some of the sequences have both SSU and LSU in the definition lines and if accurate, that should lead to an R_MultipleFamilies error. These 13 sequences that match both genes could have been trimmed before inclusion in the SilvaRef LSU set. In principle, one could detect the presence of an SSU match and an LSU match in the same sequence with RNAmmer, but one would have to add error rules to RNAmmer to decide when the occurrence of matches to both SSU and LSU models is an error. That need for error semantics exemplifies how `ribotyper` differs in functionality from RNAmmer. Of the 444 sequences that passed `ribotyper` and failed RNAmmer, 433 of them would pass RNAmmer if, as for the Yarza set, the E-value and bit score thresholds for the spotter HMMs were changed to 1 and `−100` respectively, indicating that these sequences do not match well to the eukaryotic LSU rRNA spotter profile HMM. Data and instructions for reproducing these comparisons are available at https://github.com/nawrockie/ribovore-paper-2021-supplementary-material.

In general, there are a large number of discrepant outcomes and neither RNAmmer nor `ribotyper` is consistently more restrictive than the other. We infer merely that the two pieces of software solve different problems and there is not a straightforward way to modify RNAmmer to solve the problem of checking rRNA submissions to GenBank. This helps to justify why we developed the new software Ribovore. As explained above, Ribovore also has additional modules, such as `ribodbmaker`, that are even less comparable to RNAmmer and solve other problems in rRNA sequence validation and curation.

### Limitations and future directions

Ribovore includes 18 profile models (Table 3), only two of which are used for automated submission checking (bacterial SSU rRNA and archaeal SSU rRNA), and seven of which (the first seven rows in Table 3) have been used in the context of `ribodbmaker` to generate one or more `blastn` databases or RefSeq records. Eight of the remaining models were created from alignments of fewer than 10 sequences, and need to be improved by adding more sequences. However, some of the models, especially those based on Rfam alignments such as eukaryotic SSU and LSU rRNA, could in principle be used for submission checking by `ribosensor` and we plan to investigate those possibilities based on empirical testing in the future. Beyond the existing models, more models are needed for other rRNA genes and taxonomic domains, such as mitochondrial LSU rRNA, *Microsporidia* LSU rRNA, eukaryotic 5.8S rRNA and 5S rRNA. Rfam includes alignments for some of these (e.g. 5.8S and 5S rRNA) and future versions of Ribovore could include models based on those, but manual curation effort will be required to create others.

One limitation of Ribovore is that there are many parameters and the user may need to choose the settings carefully for each distinct purpose. For example, the usage of `ribodbmaker` should be tuned for each gene and taxonomic domain, as we have reported here for fungal SSU and LSU rRNA to require the commonly targeted regions of those respective genes to be present in the sequences. Another limitation is that we do not model introns, simply expecting any introns to be un-aligned in the `ribotyper` and `riboaligner` analysis. Additionally, minimum criteria (e.g. minimum score and coverage values) for passing sequences in the `ribotyper`, `ribosensor` and `riboaligner` tools should be set based on empirical testing, and the default values for those programs are currently tailored to prokaryotic SSU rRNA based on our internal testing. Expansion to other genes and taxonomic domains will require additional testing of those values.

For some applications, the running time of Ribovore programs can be a significant limitation. Profile-based CM or profile HMM methods that compare a few profiles (in this case, at most 18) to each input sequence can be more efficient than single-sequence based methods like `blastn` which typically compare many database sequences (in this case, more than 1000) to each input sequence, but of course this depends on the relative speed of each profile to sequence and sequence to sequence comparison. CM methods that score both sequence and secondary structure conservation are computationally complex, requiring time *O*(*N*^4^) for a sequence of length N without heuristic filtering or banded dynamic programming approaches [49]. Even with heuristic-based acceleration strategies, alignment of a single full length LSU rRNA sequence typically takes several seconds on a single CPU. For this reason, `ribotyper` and `ribosensor`, which are intended to handle sequence submissions of up to millions of sequences do not, by default, compute an alignment using the CM, but rather use only more efficient profile HMM algorithms. The `riboaligner` and `ribodbmaker` programs, however, do compute alignments using CMs and so take longer per sequence, although the frequency with which these programs need to be run, at least for GenBank, is less. The `ribosensor` program, which runs both `ribotyper` and `rRNA_sensor`, which is `blastn`-based, combines both profile and single-sequence methods.

We measured the running time of `rRNA_sensor` by itself and `ribosensor` on 1000 16S sequences using 1, 2, 4, 8, and 16 processors. The `rRNA_sensor` program took 199s, 104s, 54s, 29s, and 17s, respectively; `ribosensor` took 362s, 196s, 106s, 61s, 39s, respectively. Thus, a submission of 1 million sequences, which is on the high end, would take hours to process given 16 processors on the host computer. The programs `ribotyper`, `ribosensor`, `riboaligner`, and `ribodbmaker` all include a command-line option -`p <n>` that enables parallelization by splitting the input file into `<n>` roughly equal sized chunks and processing each chunk independently on `<n>` nodes on a compute cluster. However, for `ribodbmaker`, only the `ribotyper` and `riboaligner` steps are parallelized by splitting the input file into chunks of similar size.

While doing the comparison of curated fungal datasets, we also timed `ribotyper`, `riboaligner`, and `ribodbmaker` on the 8,770 sequence Yarza fungal SSU set and the 2,993 sequence SilvaRef fungal LSU set as described in Implementation. The `ribotyper` program required 40m 14s (0.275s per sequence) on the Yarza set and 16m 51s (0.338s per sequence) on the SilvaRef set. The `riboaligner` program took 116m 22s (0.796s per sequence) on the Yarza set and 123m 56s (2.48s per sequence) on the SilvaRef set. The `ribodbmaker` program took 49m 49s wall-clock time and 1191m 2s cumulative time for all processors on the Yarza set and 60m 24s wall-clock time and 980m 11s cumulative time for the SilvaRef set. In general, we conclude that these analyses are tractable for tens of thousands of sequences at a time.

## Conclusions

Our primary contribution described herein is the software package Ribovore for rRNA sequence analysis. At NCBI since July 2018, Ribovore has been used to check the quality of incoming submissions and to curate datasets of high quality sequences for RefSeq or to use as `blastn` databases. In the submission checking context, Ribovore has been used to check nearly 50 million 16S bacterial and archaeal SSU rRNA sequences through May 31, 2020 and millions more after that date. Ribovore has also been used manually by GenBank indexers when `blastn` analyses gave uncertain results for other rRNAs. A subset of the `blastn` databases created by Ribovore are selectable by users of the BLAST webpage as target databases, and are used in over 2,000 web `blastn` runs per day. We also are using Ribovore internally to curate fungal RefSeq records for SSU and LSU rRNA from type material. We showed that this curation effort is complementary to the larger Silva effort, as it selects only the best sequences that pass a larger battery of tests. Furthermore, the RefSeq records are linked within Entrez to other NCBI resources including BioCollections, BioProjects, Taxonomy, and BLAST. With this formal report of how Ribovore is designed and implemented, we hope that both producers and consumers of rRNA sequence data will achieve a new understanding of how rRNA sequences are curated in GenBank, RefSeq, and associated resources.

## Availability and requirements

Project name: Ribovore

Project home page: https://github.com/ncbi/ribovore

Operating system(s): Linux, Mac/OSX

Programming language: Perl

Other requirements: BLAST+ v2.11.0, Infernal v1.1.4, Sequip v0.08, VecScreen plus taxonomy v0.17 (see Table 2)

License: public domain

Any restrictions to use by non-academics: none

## Abbreviations

NCBI: National Center for Biotechnology Information
rRNA: ribosomal RNA
SSU rRNA: small subunit ribosomal RNA
LSU rRNA: large subunit ribosomal RNA
CM: covariance model
HMM: hidden Markov model
nt: nucleotides
Kb: kilobase (1000 nucleotides)

## Declarations

### Ethics approval and consent to participate

Not applicable.

### Consent for publication

Not applicable.

### Availability of data and materials

All data generated or analyzed during this study are included in this published article, its supplementary material, or NCBI’s GenBank database. Code is available on GitHub (https://github.com/ncbi/ribovore). BLAST databases are available in the directory https://ftp.ncbi.nlm.nih.gov/blast/db/. The supplementary material is available on GitHub (https://github.com/nawrockie/ribovore-paper-2021-supplementary-material) and includes instructions for reproducing the comparisons reported in the article.

### Author’s contributions

AAS and EPN conceived of and designed the software and wrote most of the paper. RM, BR, CS assisted by writing passages about *Fungi* and about practical usages of Ribovore within NCBI. EPN wrote most of the Ribovore code and AAS wrote rRNA_sensor. All authors participated in the design and user interface of at least one Ribovore module. EPN, AAS, RM, BR, AJ, BAU formally tested the software. RM curated the rRNA databases. BR selected and curated the fungal RefSeqs. CS guided the multiple usages of NCBI taxonomy in Ribovore and corrected taxonomy inconsistencies as they were detected. EPN, RM, AJ, and BAU collected data on Ribovore usage. RM, AJ, and BAU used Ribovore to evaluate submissions to GenBank. IK-M supervised the work of RM, AJ, and BAU. All authors read and edited multiple versions of the manuscript and approved the final version.

## Acknowledgements

Thanks to our NCBI colleagues Alex Kotliarov and Sergiy Gotvyanskyy for assistance in integrating Ribovore into GenBank processing pipelines and for collecting data on Ribovore usage. Thanks to our NCBI colleague Richa Agarwala for providing access to an isolated Linux computer on which we could do sole-user measurements of running time.

## Figures

### Additional Files

Supplementary material available at: https://github.com/nawrockie/ribovore-paper-2021-supplementary-material.

